# Multiple contact sites between cells and the vitelline envelope coordinate tissue flows in *Drosophila* gastrulation

**DOI:** 10.64898/2026.07.30.741527

**Authors:** Marina B. Cuenca, Wan Yee Yau, Giulia Serafini, YaeJi Kim, Carl Modes, Pavel Tomančák

**Affiliations:** Max Planck Institute of Molecular Cell Biology and Genetics, 01307 Dresden, Germany; Center for Systems Biology, 01307 Dresden, Germany; Cluster of Excellence Physics of Life, Technische Universität Dresden, Dresden, Germany

## Abstract

Gastrulation is thought to be driven primarily by forces generated within individual cells. These cell-intrinsic forces collectively induce tissue-scale flows and transform the monolayered embryo into a multilayered structure. However, as the embryo constitutes a mechanically closed system, these flows must be balanced by regions of resistance or anchoring to enable asymmetric morphogenesis. In the *Drosophila* embryo, integrin-mediated attachment of the blastoderm to the vitelline envelope has been shown to stabilize germ band extension at the organismal scale. Disrupting such an attachment leads to a characteristic twisting phenotype. Yet, how this attachment shapes concurrent global morphogenetic events remains unclear. We discovered that the integrin *α*-subunit scab, which mediates the attachment, is expressed in three different regions of the cellular blastoderm near prominent invagination events. Through a combination of light-sheet imaging, genetic and mechanical perturbations, we demonstrate that integrin-enhanced friction is essential for unidirectional tissue flows in those regions, with effects including cephalic furrow positioning and epithelial stability. Guided by a minimal physical model, we further show that multiple attachment sites enhance the robustness and reproducibility of global tissue movements. Together, our results indicate that *Drosophila* gastrulation emerges from a balance between cell-intrinsic force generation and spatially distributed adhesion to the surrounding envelope, which together shape tissue flows at the embryo scale.

## 1. Introduction

Epithelial tissues undergoing morphogenesis can be described as active viscoelastic surfaces, in which tissue-scale flows emerge from the balance of internally generated stresses and external constraints [1]. In such systems, large-scale deformation modes like drift and rotation are generically present and can lead to variability in global flow patterns. Theoretical descriptions of active epithelial sheets and shells have shown that these modes are strongly influenced by boundary conditions and geometric constraints [2]. For example, during wing disk morphogenesis in the *Drosophila* larval stages, attachment of this tissue to the surrounding ECM has been shown to guide proper folding [3]. In parallel, experimental work has demonstrated that adhesion of the blastoderm to the vitelline envelope, the egg-shell innermost layer, mediated by integrins provides external forces that counteract tissue-intrinsic contractility and shape gastrulation movements [4, 5, 6, 7, 8].

In *Drosophila melanogaster* and *Tribolium castaneum*, attachment occurs in the vicinity of major invagination events, although at different positions along the embryo surface. In *Drosophila*, the attachment is mediated by integrin *α*-subunit scab which is expressed anterior to the posterior midgut (PMG) invagination site [9]. Loss of this integrin can lead to a characteristic twisting of the embryo during germ band (GB) extension, an inherently unstable process [5, 10, 8]. Similar perturbations in *Tribolium* result in a symmetric pattern of tissue flow, suggesting that the attachment contributes to the global coordination of morphogenesis and may act as a mechanical constraint that breaks symmetry during gastrulation resulting in unidirectional tissue flows [5].

Although organism-level consequences of integrin-mediated attachment have been described, its impact on tissue dynamics at smaller scales remains only partially understood in *Drosophila*. Previous work has analyzed tissue flows in the vicinity of the PMG attachment site, providing insight into how local attachment influences immediate tissue movements [6, 7]. However, it remains unclear how the absence of attachment affects tissue flows across the embryo as a whole. In particular, whether and how local perturbations propagate to distant regions, and how multiple morphogenetic events are coordinated through these effects, has not been quantitatively assessed. Addressing this requires resolving tissue movements with sufficient spatial and temporal resolution to capture both local flow patterns and their integration at the whole-embryo level.

In this study, we show that integrin-mediated attachment of the blastoderm to the vitelline envelope occurs at three spatially distributed sites in *Drosophila*: near the posterior midgut invagination, the dorsal cephalic furrow (CF), and the ventral CF. All three are required for coordinated tissue flows during gastrulation. Using light-sheet imaging and particle image velocimetry, we demonstrate that loss of the integrin *α*-subunit scab disrupts tissue flow in all three regions within a defined 10–20 minute window of GB extension, with downstream consequences including impaired CF displacement and ectopic epithelial buckling. A minimal physical model of the embryo cross-section confirms that attachment sites not only act locally but also influence morphogenetic landmarks at a distance, and that three spatially separated attachments are sufficient to enhance the reproducibility of tissue movements at the embryo scale.

## 2. Results

### 2.1. Integrin-mediated attachment breaks symmetry in tissue flows in morphogenesis

To understand this, we used *Drosophila* as a model system to analyze the tissue flows at the cellular level in scab loss-of-function embryos compared to the well-established wild-type (wt) behavior. Following a previously used approach for *Tribolium* [5], we imaged *Drosophila* wt and *scab* mutant embryos during gastrulation using light-sheet microscopy (Fig. 1a, a’ and Video S1) [11]. Since this technique allows for optical tissue sectioning, we used it to record the complete sagittal cross-sections of the embryo with temporal resolution sufficient to resolve tissue flows at low phototoxicity (see Methods). To increase throughput, typically limited to a single embryo, we employed an alternative mounting method that glued samples to a glass capillary in the desired orientation [12]. This method enabled imaging of up to 10 samples with 15-second time steps. Recordings were temporally aligned, with time 0 corresponding to the onset of gastrulation, identified by the apical constrictions leading to the ventral furrow (VF) invagination.

**Figure 1.**
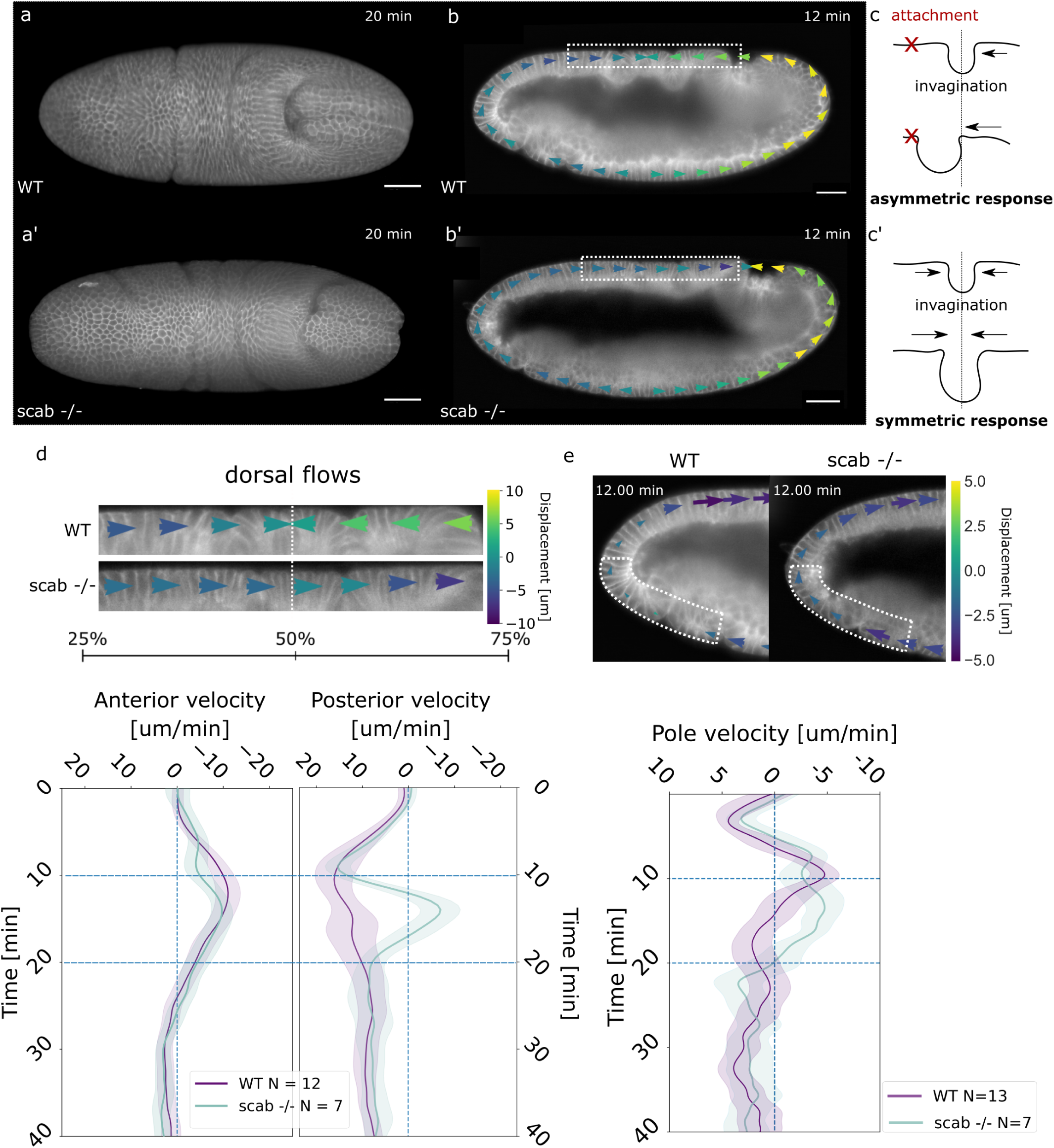
Integrin-mediated attachment breaks symmetry in tissue flows. (a, a’) Dorsal view 3D rendering of *Drosophila* wt and scab loss-of-function embryos during gastrulation. (b, b’) Sagittal section and quiver plot of displacement field quantified with PIV analysis in wt and *scab* mutant. (c, c’) Diagram of symmetry breaking during invagination in the presence of an attachment and symmetric flows in the absence of attachment. (d) Zoomed-in region of dorsal displacement field in wt and *scab* mutant, quantified averaged velocities and 95% CI on the anterior (25–50% EL) and posterior regions (50–75% EL). (e) Zoomed-in region of ventral-anterior displacement field in wt and *scab* mutant and quantified average velocity and 95% CI in the portion indicated in dotted line area. Dotted blue lines in the plots indicate the time window where tissue flows are altered. Scale bar, 50 *µ*m.

To quantify tissue flows in the 2D cross-sections, we focused on wt and *scab* mutant embryos without the strong twisting phenotype, as this would confound the 2D flow analysis. Preliminary observations and kymographs of the dorsal tissue showed a wide variability in GB extension dynamics in integrin mutants (Fig. 1a, a’ and Fig. S1a, b, c). To quantify these tissue flows, we used particle image velocimetry (PIV) to quantify local tissue velocity from the 1D-curved surface of the blastoderm during GB extension, and categorized the direction of the flows as clockwise (negative velocity values) and counterclockwise (positive velocity values), orienting the embryos with the anterior side to the left. In wt embryos, the dorsal tissue exhibited confluent asymmetric tissue flows towards the center of the embryo while *scab* mutants showed a unidirectional positive displacement towards the PMG invagination, consistent with a lack of attachment of this tissue and previously reported results (Fig. 1b, c) [6].

To quantitatively compare these differences, we performed averaging of the velocities in the anterior and posterior halves of the dorsal tissue. We observed that the posterior half, near the PMG invagination, exhibited a dramatic direction reversal from positive to negative velocity values between 10–20 minutes of GB extension in *scab* mutants (Fig. 1d, Video S2). This was consistent with the qualitative observation of counterflows in kymographs in the same region (Fig. S1a, b, c). Without attachment, symmetric tissue flow emerges (Fig. 1c, c’), similar to observations in *Tribolium* embryos [5] or during mesoderm invagination [13]. This flow pattern supports scab’s role as an anchor to the vitelline envelope, where attachment ensures asymmetric tissue responses.

Next, we investigated if the patterns of tissue flow were affected elsewhere around the circumference of the embryo upon lack of attachment. We noticed that, in the anterior part of the dorsal region, away from the known attachment, the *scab* mutant embryos showed impaired clockwise tissue flow during the early minutes of GB extension; subsequent displacement was similar to wt embryos (Fig. 1d). This suggests a possible long-range influence of scab activity. The PIV analysis also identified altered tissue flow in *scab* mutants in a remote region of the embryo in the ventral portion of the anterior pole, far away from the known attachment. Initially, both wt and *scab* mutants followed the same flow pattern, switching from counterclockwise to clockwise (Fig. S1d, e). At 10–20 minutes of GB extension, wt flow stopped, whereas *scab* mutants continued to exhibit significant clockwise cellular flow (Fig. 1e; further Fig. S1f, g). Taken together, the PIV analysis of *scab* mutants revealed three regions with altered tissue flows (Fig. 2a), only one of which is proximal to a reported attachment to the vitelline envelope.

**Figure 2.**
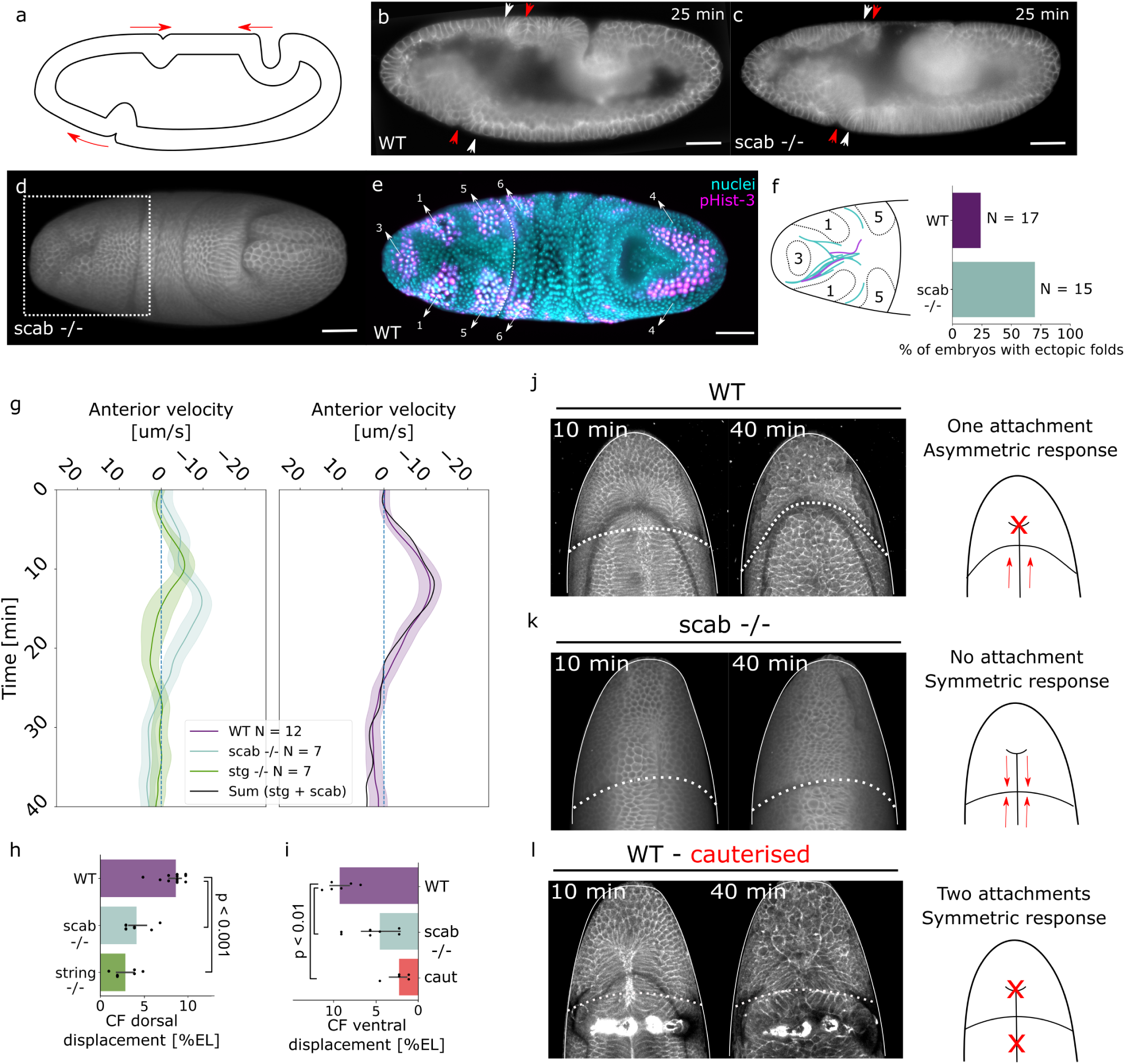
Attachment affects cephalic furrow positioning and epithelial stability. (a) Diagram of affected tissue flows in the embryo surface relative to invaginations. (b, c) Sagittal sections of wt and *scab* mutant embryos highlighting CF displacement. White arrows show initial position at the start of invagination and red arrows show final location. (d) Dorsal 3D rendering of scab loss-of-function with dotted white box highlighting ectopic folds in the head region. (e) Dorsal 3D rendering of wt embryo stained for nuclei and pHist-3 showing mitotic domain regions, numbered according to division order. (f) Diagram of mitotic domains, location of ectopic folding and frequency of the folds in wt and *scab* mutant embryos. (g) Average velocities and 95% CI of the anterior dorsal region in *scab* and *stg* mutants, plus the comparison of adding up these two curves recapitulating the behavior in the wt. (h) Barplots of average, 95% CI and single data points of quantified CF dorsal displacement in wt, *scab* and *stg* mutants. (i) Barplots of average, 95% CI and single data points of quantified CF ventral displacement in wt, *scab* mutant and cauterized wt. (j, k, l) Ventral 3D rendering of head region in wt, *scab* mutant and wt cauterized embryos showing CF shape in dotted white line and diagram of the underlying attachment hypothesis. Note: (j, l) present a more pronounced inverted U-shaped shadow from the contact area between the tissue and the coverslip, not to be confused with the cephalic furrow. Scale bar, 50 *µ*m.

### 2.2. Integrin-mediated attachment affects cephalic furrow positioning and epithelial stability

The two altered tissue flow patterns at the anterior end of the embryo occurred in the vicinity of the cephalic furrow (CF). The CF is a highly stereotypical morphogenetic event initiated by the lateral shortening of a single row of blastoderm cells positioned at the future head-trunk boundary by a well-understood gene regulatory network [14, 15]. To determine whether the altered tissue flows impact CF formation, we examined the dynamics of this morphogenetic event in the *scab* mutant. The CF invagination begins laterally and propagates both ventrally and dorsally, forming a closed ring around the embryo’s surface [16]. In the wt embryo, as the tissue internalizes, the CF angle relative to the dorsoventral axis increases, producing a posterior shift of the dorsal and anterior shift of the ventral regions as if the furrow were, as a whole, rotating clockwise [15, 16]. In *scab* mutant embryos, due to changes in the flow pattern of the blastoderm, this CF movement is impaired (Fig. 2b, c, h, i). Moreover, immediately anterior to the dorsal edge of the furrow, in the head region of the embryo, we observed frequent Y-shaped buckling of the blastoderm tissue (Fig. 2d, Video S3). We also observed this characteristic buckling of the epithelium in wt embryos, albeit with a much lower frequency (Fig. 2f).

Recent studies of CF function in *Drosophila* revealed that abrupt ectopic buckling of the blastoderm epithelium is caused by compressive stresses originating from local apical area expansion of mitotic domains and global push of the GB in the head-trunk boundary region [15, 17]. In the context of the *scab* mutant, we hypothesized that a similar situation occurs. Within the tissue flow divergency time window (10–20 minutes after gastrulation onset), several mitotic domains in the dorsal head region undergo apical expansion, and buckling occurs between them (Fig. 2e). Moreover, the CF’s diminished displacement posteriorly resembles the GB’s global compressive effect at the head-trunk boundary.

To evaluate the relative contribution of mitotic domain expansion to the observed CF displacement, we quantified tissue flows in *string* (*stg*, *CDC25*) mutant embryos, which lack mitotic domains but otherwise do not disrupt gastrulation morphogenesis (Video S4). At the 10–20 minute interval, the *stg* mutants show a complete halt in the dorsal-anterior clockwise tissue flow (Fig. 2g), an intermediate level of CF displacement compared to wt and *scab* mutants (Fig. 2h) and, as expected, no tissue buckling. Interestingly, adding up the aligned flow values of *stg* and *scab* mutant embryos recapitulates the wt flow curve (Fig. 2g). Thus, this analysis allowed us to deconstruct the wt dorsal-anterior tissue flow into two distinct components: a cell proliferation-driven component and a scab-mediated contribution. We conclude that the lack of dorsal CF shift and the occurrence of ectopic head folds are phenotypes associated with the absence of scab expression, the latter being a consequence of concurrent compressive forces from the former and the expansion of the mitotic domains.

To investigate the most distant ventral region of the embryo affected in *scab* mutants, we analyzed the dynamics of the ventral CF shift. In wt embryos, the CF closes around the ventral side of the embryo and, as the anterior shift progresses, the profile acquires a distinct U-shape (Fig. 2j). By contrast, in *scab* mutants, the furrow remains straight (Fig. 2k). Analogous to the attachment near the PMG invagination site, this behavior in the wt is an asymmetric tissue response that suggests the presence of an attachment site anterior to the ventral CF. To test this hypothesis, we neutralized the effects of such an attachment by inducing artificial adhesion of the cells to the vitelline envelope posterior of the CF using laser cauterization. Supporting our hypothesis, cauterized embryos exhibited a phenotype similar to *scab* mutants, with the ventral CF showing minimal or no shift and maintaining a straight configuration throughout gastrulation (Fig. 2i, l). This intervention restored symmetry in tissue flows, consistent with a hypothetical attachment site at the ventral anterior pole.

### 2.3. Three distinct integrin-mediated attachments drive asymmetric flows

Having seen the global effects of scab loss-of-function around the embryo circumference away from the known scab expression domain, we hypothesized that *scab* may be expressed at additional sites on the embryo’s surface and function as an attachment or enhanced friction region. To test this, we employed the hybridization chain reaction (HCR, Molecular Instruments) protocol to visualise *scab* mRNA expression with greater sensitivity. The results revealed *scab* signal in three distinct regions in wt embryos, matching the altered tissue flows (Fig. 3a, b). At stage 5, expression was detected at the previously reported posterior-dorsal attachment site, as well as in the anterior-ventral pole. While expression at the posterior-dorsal site increased throughout gastrulation, the newly identified anterior signal diminished and became concentrated in the ventral portion of the anterior pole (Fig. 3a, b and Fig. S2). Interestingly, this newly found expression site is analogous to the positioning of the attachment previously reported in *Tribolium* [18, 5]. Additionally, low levels of *scab* expression were detected in the dorsal-anterior region near the CF at stage 6, persisting throughout gastrulation (Fig. 3a, b). Previously, in situ hybridization studies identified this site only at stage 8 [9]. Whether this region can function as an attachment has not yet been assessed. Our results indicate that the *scab* expression sites are consistently associated with areas of divergent tissue flows in gastrulating embryos detached from the vitelline envelope. From now on, we assign each attachment the identifier of its closest invagination site: **PMG**, **CFV** (CF ventral) and **CFD** (CF dorsal).

**Figure 3.**
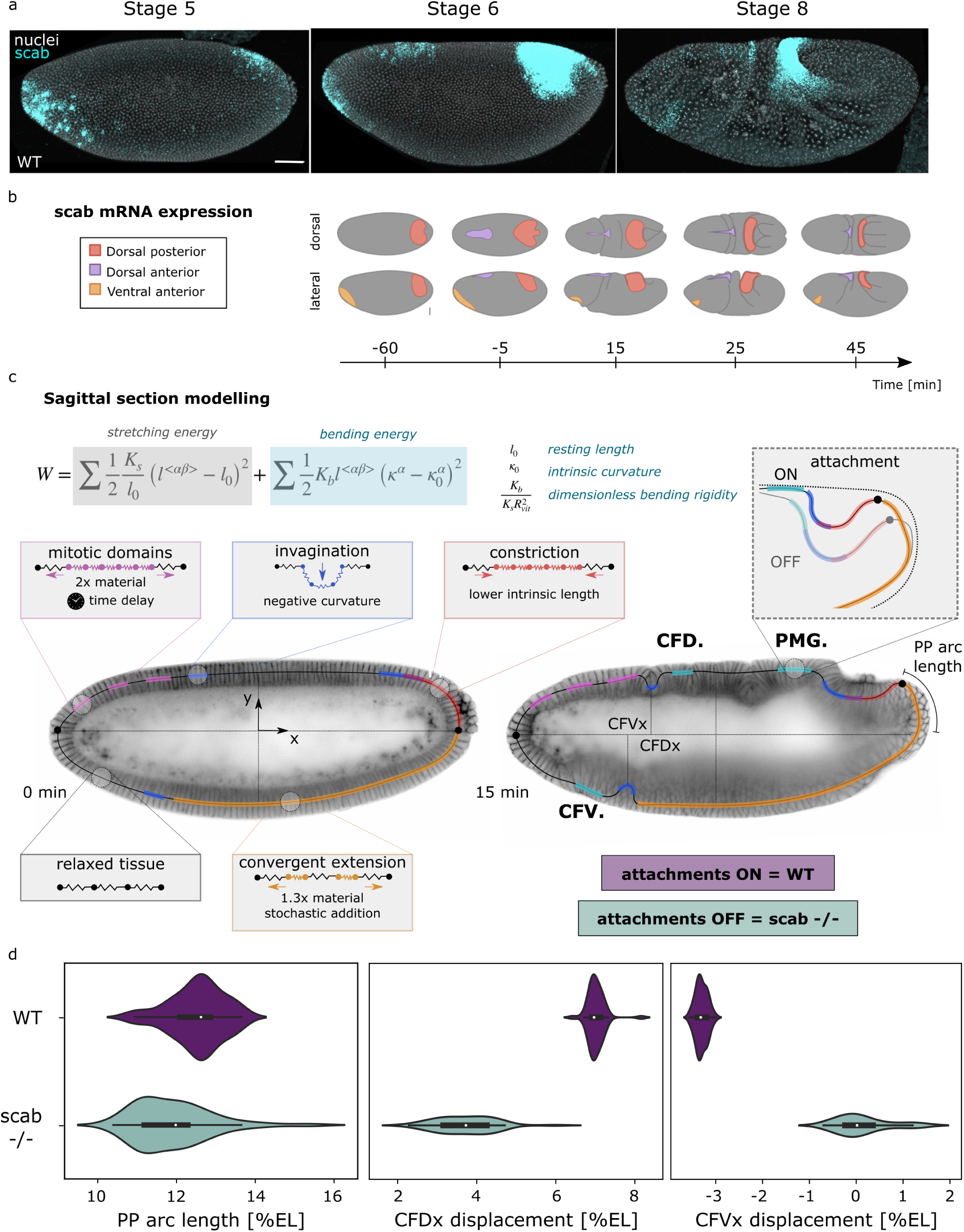
Three distinct integrin-mediated attachments and their physical modeling. (a) Maximum projection of lateral view of nuclei staining and *scab*HCR in wt embryos showing 3 distinct expression sites. (b) *Scab*mRNA expression diagram for each of the attachment sites. (c) Sagittal section modelling diagram with total energy expression and the different regions along the embryo circumference: mitotic domains, invaginations, constriction, convergent extension and relaxed tissue. Attachments are highlighted and can be turned on by fixing the position of these particles during the simulations. Morphogenetic landmarks are defined as follows: to measure GB extension, the posterior pole (PP) arc-length traveled by the tip of the GB was quantified; for the cephalic furrow, the displacement in the x-axis is quantified for the dorsal (CFDx) and ventral (CFVx) regions. All measurements are percentual relative to the total EL. (d) Distributions of the final position or displacement of relevant features in attachments ON (wt scenario) and OFF (*scab* mutant scenario). Scale bar, 50 *µ*m.

### 2.4. Minimal physical model highlights attachment role in morphogenetic robustness

To understand the impact of the attachment sites on tissue flows and morphological landmark placement, we developed a simple 2D physical model of the embryo’s circumference. The model, adapted and extended from Vellutini et al. [15] (see Methods), represents the sagittal cross-section of the stage 6 embryo as a curved, one-dimensional chain of massless particles connected by strings. The tissue is constrained by an effective vitelline envelope, an outer elliptical boundary which prevents the particles from crossing it (this boundary is not shown in the simulation representations). Tissue energy is defined by a combination of stretch and bending components, with *l*_0_ representing the spring rest length, *K_b_* the dimensionless bending rigidity and *κ* and *κ*_0_ the current and preferred curvatures, respectively (Fig. 3c).

The model incorporates regions with differential properties to simulate morphogenetic processes. Invaginating regions, such as the PMG, ventral CF and dorsal CF, are represented by local changes in preferred curvature, inducing tissue bending. In the PMG, additional tissue contraction is simulated by modifying the resting length of the springs locally. With the aim of simulating the time window spanning the tissue flow divergencies between the wt and *scab* mutants, the first 25 minutes after gastrulation onset, GB convergent extension is modeled by incrementally adding 30% more particles. Each of these particles is added with an associated new spring of rest length *l*_0_ at random positions along the ventral-posterior regions of the tissue. When 70% of these particles are introduced, mitotic domains in the head region are activated, reflecting the observed delay between these events. In these domains, particle addition is doubled and happens simultaneously. The simulation concludes when the GB finishes its displacement and the tissue reaches equilibrium.

To simulate scab-mediated adhesion to the vitelline envelope, three attachment sites were incorporated near the invagination regions (Fig. 3c), forcing particles to remain static. A total of 50 simulations were performed with and without attachments, monitoring morphogenetic landmarks to evaluate the effect of attachment presence (ON = wt) or absence (OFF = *scab* mutant) (Video S5 and S6). Consistently, parameters such as posterior pole arc length, ventral and dorsal CF displacement exhibited significantly shifted mean values and reduced robustness (higher variance) in the absence of attachments (OFF) (Fig. 3d). This resulted in altered positioning of morphogenetic events, consistent with experimental observations. Additional parameters, such as invagination depth and anterior pole displacement, showed similar trends (Fig. S3). Together, these findings underscore the critical role of scab-mediated attachments in ensuring specific, reproducible landmark placement in the developing embryo.

### 2.5. Multiple attachment sites stabilize the embryo against twist

Simulations support the idea that gastrulation morphogenesis is stabilized by three attachment sites around the embryo’s circumference, and experiments suggest that there may be up to three such sites. We next asked how that relates to the organismal-level twist observed in *scab* mutants. This twist phenotype could be expected to depend strongly on fixed attachments, particularly since three attachments are the minimal number needed to prevent global rotations and their absence would therefore be subsequently permissive to twist. Such a global twisting phenotype cannot be captured in a 2D model because the dimensionality is too low, so we return to experimental interrogation of the system.

To determine whether the quantified discrepancies between wt and *scab* mutant embryos contribute to the twisting phenotype and to assess the involvement of the newly identified attachment sites, we analyzed tissue flows in twisting *scab* mutant specimens. By acquiring cross-sections at the embryo’s equator, we performed PIV analysis focusing on the anterior pole (Fig. 4a). In both symmetric wt and *scab* mutants (non-twisting), the average velocities were close to zero, whereas in twisting *scab* mutants, cells exhibited displacement towards either side with distinct dynamics (Fig. 4b, Fig. S4a, b, c). These findings support recent observations on the twisting phenotype leading to head rotation, which is facilitated by the absence of scab-mediated attachment [8].

**Figure 4.**
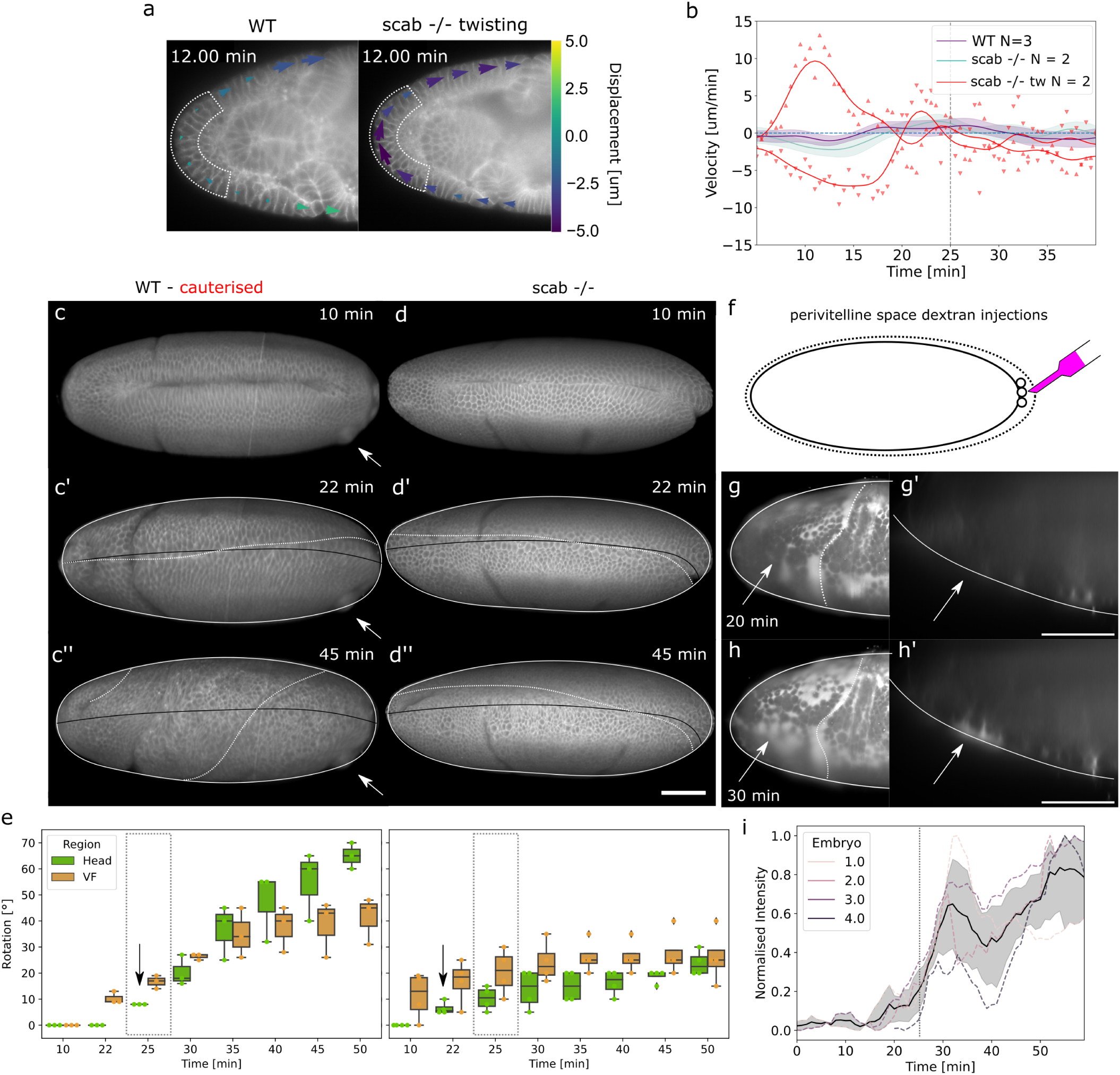
Multiple attachment sites stabilize the embryo against twist. (a) Equator section of the head with quiver plot of displacement field in wt and *scab* mutant that twists; dotted white lines indicate the region quantified. (b) Averaged velocities and 95% CI in wt, non-twisting *scab* mutant and two different directions of twisting *scab* mutants. (c, d’) Ventral 3D rendering of cauterized wt and twisting *scab* mutant embryos showing cauterized area (white arrow), initial VF position (black line), current VF position (dotted white line). (e) Boxplot of quantified VF deviation and head rotation for cauterized wt (*N* = 3) and twisting *scab* embryos (*N* = 4) highlighting first detection of head rotation (black arrow) and mitotic divisions in domain 8 (gray dotted box). The box represents the quartiles and error bars the full distribution minus outliers. (f) Diagram of perivitelline dextran injections performed between the end of cellularization and gastrulation onset. (g, g’) Ventral 3D rendering and sagittal section of the head region before mitotic domain 8 division in wt showing dextran exclusion suggesting cells are attached to the vitelline envelope. (h, h’) Ventral 3D rendering and sagittal section of the head region after mitotic domain 8 division in wt showing dextran signal suggesting cells are detached from the vitelline envelope. (i) Average dextran intensity, 95% CI and single curves showing signal increase upon mitotic domain 8 division (gray dotted line).

To characterize this head rotation, we designed an experiment to induce the twisting phenotype in wt embryos through laser cauterization. Specifically, an artificial attachment was generated between the posterior lateral cells and the vitelline envelope to create an artificial permanent anchor leading to asymmetric tissue flows. The ventral portion of the embryo was then imaged to quantify head rotation angles and VF deviation (associated with GB twisting). In all three cauterized wt embryos, a pronounced twist of the embryo was observed, which was then compared to twisting *scab* mutants (Fig. 4c, d, Video S7). In all twisting embryos, VF deviation preceded head rotation, with the latter counteracting the direction of the former (Fig. 4e, Suppl. Fig. 5). However, *scab* mutant embryos exhibited an earlier onset of head rotation compared to cauterized wt embryos (Fig. 4e). This is consistent with the lack of attachment during this time window.

The dynamics of the ventral attachment site could be related to cell cycle regulation given that cells disassemble their adhesive structures during division. We investigated whether cells in the anterior ventral pole detach in coordination with division within mitotic domain 8 [19], which overlaps with the ventral attachment site (Fig. S4d-f). To test this hypothesis, we injected fluorescent dextran into the perivitelline space of wt embryos immediately after cellularization and imaged the ventral region (Fig. 4f). The results showed dextran exclusion from the anterior ventral pole during the initial minutes of gastrulation (0–20 min), suggesting close cell contact with the vitelline envelope (Fig. 4g, h). The subsequent increase in dextran signal coincided with the division timing of mitotic domain 8 (Fig. 4i). These findings indicate a potential transient function of the newly identified ventral *scab* expression site, which acts as a stabilizing mechanism during rapid GB extension to prevent left-right asymmetries from propagating across the embryo’s surface. We hypothesize that the torque generated by the GB during the twist induces a compensatory counter-rotation of the head. In the wt condition, minor deviations during early GB extension can be corrected due to the CFV attachment. Larger perturbations induced by cauterization or by the global loss of attachment ultimately result in loss of left-right symmetry.

### 2.6. Minimal physical model suggests influence of attachment sites on remote embryo regions

A question that remains unresolved is how individual attachment sites influence tissue flows in the embryo. Experimentally probing these different states is unfortunately currently not possible. Our attempts to mimic combinations of attachment sites in *scab* mutants using laser cauterization do not accurately replicate the dynamics of integrin-mediated attachments. Furthermore, selectively abolishing existing attachments in wt embryos proved to be mechanically unfeasible with techniques such as oil and ferromagnetic oil injections, and no genetic approaches to selectively interfere with individual attachments are currently available. Optogenetic RNA interference experiments were performed with the ShineGAL4/UAS system [20], but did not effectively induce Scab loss-of-function phenotypes at the gastrulation stage. Our minimal theoretical model, however, is capable of interrogating different combinations of attachment sites being on or off.

To that end, we simulated scenarios with only one or two attachments and compared the positions of the selected landmarks with those from the fully ON/OFF simulations (Fig. S5). To simplify the interpretation of the results, simulations with one attachment (denoted *scab* + N, where N = PMG, CFD, CFV are the indices of the three attachment sites which could be individually turned on, Fig. 5a) and two attachments (denoted wt *−* N similarly, except here the index N is for the attachment site that is turned off, Fig. 5a’) were plotted and analyzed independently (Sup. Fig. 4c, d). The results were ranked between the *scab* (all OFF) and wt (all ON) distributions to assess the relative contribution of each attachment to specific landmarks.

**Figure 5.**
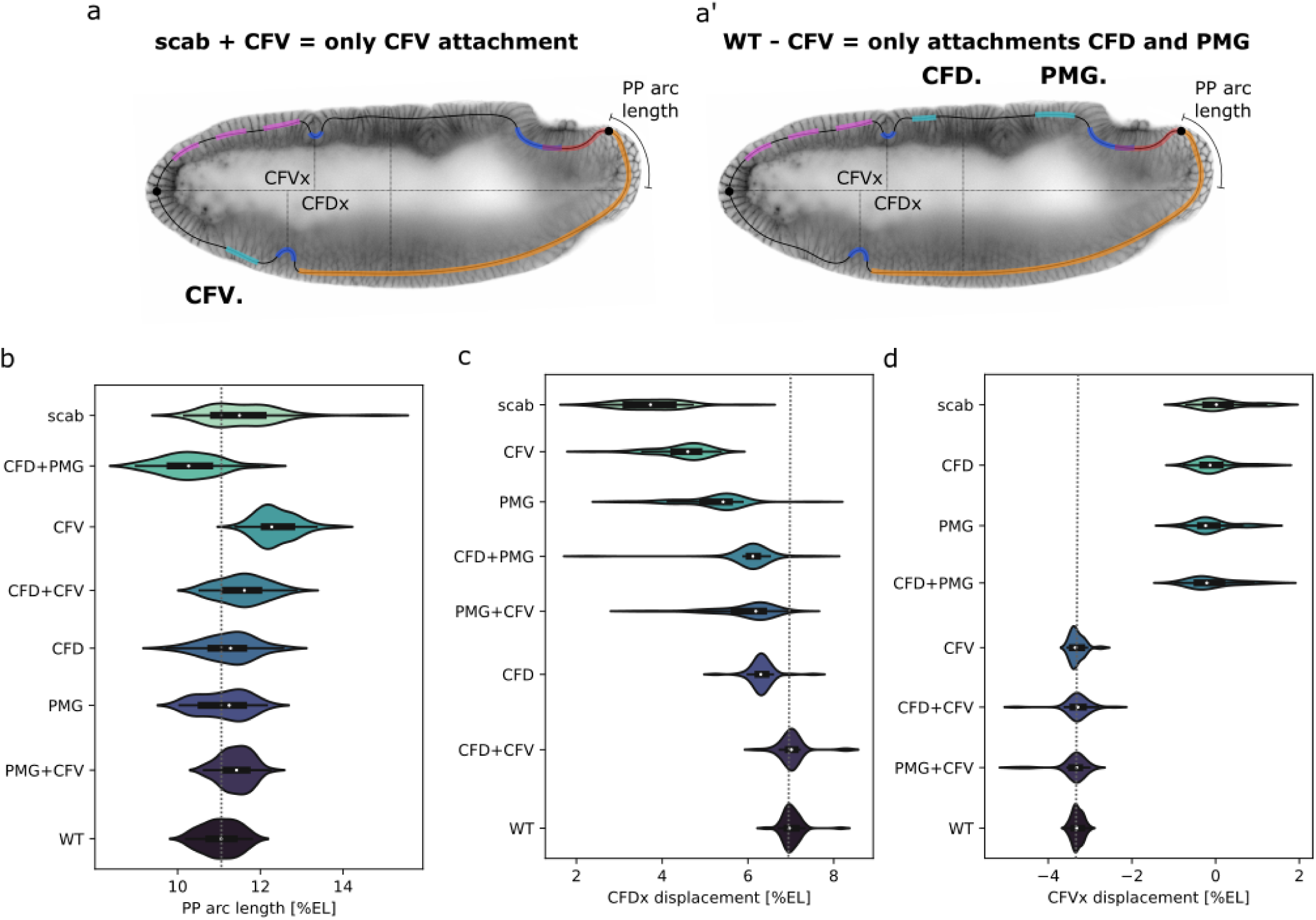
Single and double attachment simulations reveal remote mechanical coupling. (a, a’) Diagram and notation for simulations with one and two attachments. (b, c, d) Violin plots of the distribution of quantified feature locations ranked by relevance in the single and double attachment situations compared to all attachments on (wt) and all OFF (*scab*). The gray dotted line highlights the mean of the wt situation.

Initial observations revealed three distinct patterns. First, some landmarks exhibited overshooting contributions of some attachments that were compensated by others, such as the PP arc length and PMG depth. The posterior pole arc length exhibited a continuous shift in distribution but showed a characteristic overshooting associated with the CFV attachment. Specifically, when only the PMG or CFD attachments were present, the arc length showed intermediate mean values with high standard deviation (Fig. 5b). However, having only the CFV attachment produced a distribution with slightly higher values than the wt scenario. This pattern was consistent in two-attachment simulations, where the combination of the PMG attachment with either CFV or CFD resulted in elevated mean values compared to wt (Fig. 5b). Notably, when all attachments were present except for the PMG, the distribution was less robust than that observed in *scab* mutants (Fig. 5b). Similarly, the PMG depth exhibited a comparable pattern for its closest attachment (Sup. Fig. 6b). These findings underscore the importance of attachments PMG and CFV in generating robust landmark configurations for PMG depth and posterior pole arc length, associated with GB extension.

Then, some landmarks had distributions that fell between the mean *scab* and wt, such as the dorsal CF displacement and depth. While all attachments contributed individually, the CFD attachment alone produced results closest to wt (Fig. 5c, Sup. Fig. 6c). Finally, landmarks associated with the CFV attachment, including anterior pole displacement (Sup. Fig. 6a), ventral CF displacement (Fig. 5d) and depth (Sup. Fig. 6c), demonstrated a binary dependence on this attachment site. These results underscore the potential significance of the newly found CFV attachment not only in establishing robustness at the ventral anterior region of the embryo but also in contributing to the GB extension process, as reflected by changes in the posterior pole arc length and PMG depth. A previous study discussed the CF as a whole as a structure providing long-range mechanical guidance to GB extension [21]. Our work adds a layer to the CF dynamics shaping those long-range interactions.

## 3. Discussion

In this study, we dissected the disruption of tissue flows resulting from the absence of integrin-mediated attachment and identified additional phenotypic differences in CF dynamics and epithelial stability compared to wt embryos. These differences originated from previously unreported local attachment sites near prominent invagination events, revealing a total of three critical regions where scab is required for proper gastrulation. Disruption of scab-mediated attachment altered tissue flows in all three regions within a defined time window, between 10 and 20 minutes after gastrulation onset.

Notably, this time window coincides with the phase of rapid germ band extension, a process known to be mechanically unstable and highly sensitive to perturbations [10, 8]. During this phase, large-scale tissue movements generate significant shear and can give rise to embryo-scale torque. In the absence of sufficient mechanical constraints, such as integrin-mediated attachment, these forces can be accommodated through global modes of deformation, including rotational flows, ultimately leading to the characteristic twisting phenotype. This suggests that attachment sites are particularly critical during this transient regime, where they act to buffer mechanically unstable dynamics and prevent the amplification of asymmetries into organism-scale deformations.

From a mechanical perspective, multiple blastoderm– vitelline attachments are expected to enhance robustness because they act as discrete constraints on an active curved epithelial shell. Whereas one or two symmetric attachments along the AP axis still permit embryo-scale sliding and long-wavelength rotational modes, three spatially separated contact sites are sufficient to define a stable geometric reference frame, suppress soft global flow modes of the viscoelastic material, and partition the tissue into mechanically coupled domains. In this way, the resulting superficial flows become less sensitive to local fluctuations in active stress or material properties and therefore more reproducible at the whole-embryo scale. Our observations are consistent with this interpretation, suggesting that spatially distributed attachment points restrict the space of admissible flow fields and thereby stabilize the embryo-scale morphogenetic program [1, 5, 2].

To better understand these dynamics, we developed a physical model representing the sagittal section of the embryo and incorporating key morphogenetic events, including mitotic divisions. Simulations with and without attachments showed that the absence of attachments led to less reproducible landmark positioning, consistent with the experimentally observed defects in CF displacement and GB extension. However, because the model treats the embryo as a one-dimensional curved surface, it cannot capture symmetry-breaking events such as twisting. We used the physical model to explore combinations of one or two attachment sites during gastrulation, a scenario not currently accessible experimentally. While certain landmarks, such as ventral CF displacement, depended primarily on the local attachment, most landmarks were also influenced by remote attachment states. Notably, the ventral anterior attachment had a substantial effect on GB extension, highlighting that attachment sites contribute not only locally but also through long-range mechanical coupling across the embryo surface.

An additional point raised by our data is the evolutionary context of the newly identified ventral anterior attachment. In *Tribolium castaneum*, integrin-mediated adhesion of the blastoderm to the vitelline envelope was reported in an anterior-ventral domain and proposed to bias tissue flow toward a unidirectional mode during gastrulation [5]. Our finding of a ventral-anterior scab-positive domain in *Drosophila melanogaster* therefore does not appear as an isolated novelty, but rather as a related mechanical solution deployed in a distinct embryonic context. Although the spatial relationship between extra-embryonic tissues and invagination events differs substantially between short-germ and long-germ insects [22, 23], both systems appear to use localized blastoderm–envelope attachment to impose directional bias and suppress symmetric or misdirected tissue movements during gastrulation [5].

In this light, it is also not entirely unexpected that attachment-associated *scab* expression is found near both posterior and anterior gut-associated morphogenetic domains. In *Drosophila*, foregut and hindgut formation arise from anterior and posterior invagination events, and terminal patterning genes such as *fork-head*, *caudal*, and *brachyenteron* are central to gut specification and morphogenesis [24, 25]. Comparative work in *Tribolium* has further shown that major elements of terminal and gut patterning are conserved across insects, including *fork-head* expression at the termini and a conserved role for *brachyenteron* in posterior gut development [26, 27]. We therefore speculate that the recurrent association of attachment sites with gut-associated regions may reflect an evolutionarily conserved coupling between terminal and gut patterning programs and local mechanical interactions with the vitelline envelope.

In conclusion, our results indicate that all three attachment sites contribute to the coordination of tissue flows during gastrulation at both local and organismal scales. Together, they support GB extension, promote the proper dorsal shift of the CF to prevent ectopic buckling, and stabilize the anterior region against head counter-rotation arising from defects in early GB extension.

## 4. Materials and Methods

### 4.1. Fly stock keeping and transgenic lines

To generate fluorescent integrin and mitotic domains mutants, we performed genetic crosses using the loss-of-function alleles *stg* 2 (FBal0247234) and *scb*2 (FBst0003098); the membrane fluorescent marker*Gap43::mCherry* (FBal0258719, gift from Kassiani Skouloudaki); and the green fluorescent balancers *FM7c, Kr−GFP* (FBst0005193), *CyO, twi−GFP* (gift from Akanksha Jain), and *TM3, Kr−GFP* (FBst0005195).

For *stg*, which is located the III chromosome, we recombined the allele with the *Gap43::mCherry* marker, obtaining the line *Gap43::mCherry*, *stg2/TM3, Kr−GFP*. *Stg2* homozygous embryos were identified by the lack of cell divisions after gastrulation. For *scab*, we employed the line *scb2/CyO, twi-GFP; RGAPm::Cherry/RGAP::mCherry* [8], where the *RGAP::mCherry* is recombined and viable in homozygosis. *Scb2* homozygous embryos were identified by absence of twi-GFP fluorescence after imaging.

Stocks were kept and embryos were collected as described in [15].

### 4.2. HCR in situ hybridization

Hybridization Chain Reaction in situ hybridizations were performed according to previous studies [28, 8] with the following modifications. Fixed embryos conserved in 100% MeOH at *−*20*^◦^*C for at least one day were rehydrated and incubated for 7 min at room temperature in 1 mL of 10 *µ*g/mL proteinase K (Cat. No. P2308, Sigma-Aldrich) solution in PBT 0.1%. After the post-fixation step in 4% PFA in PBT 0.1%, the samples were rinsed 6 times in 1 mL PBT 0.1%. Incubation with the RNA probes lasted 21 hours, while the incubation with the hairpin solution was usually 19 hours. After the washes to remove the hairpin solution, DAPI staining was performed in a 1:1000 20 mg/mL DAPI in 5*×* SSCT solution for 2 h at room temperature. If not imaged the next day, prepared samples were kept in 60 *µ*L SlowFade Diamond mounting medium (Thermo Fisher Scientific) at 4*^◦^*C for up to one week, to be mounted only on the same day or the day before imaging.

Samples were imaged by laser-scanning confocal microscopy (Zeiss LSM 880 Airy upright, equipped with two PMT and one GaAsP detector) using the LD LCI Plan Apo 25*×*/0.8 immersion objective (Zeiss) in glycerol immersion, with a zoom factor of 0.6*×*. Serial sections with a Z-resolution of 2 *µ*m covering approximately 60% of the embryo’s volume were acquired with Zeiss ZEN 2.3 SP1 FP3 (black version) software. Laser lines used were 405 nm (DAPI nuclei), 561 nm (AlexaFluor-546) and 633 nm (AlexaFluor-647). Pinhole size was 40 *µ*m, pixel dwell time was 2.05 *µ*s, pixel size was 0.554*×*0.554*×*1.452 *µ*m. For *scab* mRNA signal intensity quantification, the 561 nm laser power used was 0.252 mW. Images were processed using a custom ImageJ macro in Fiji [29].

### 4.3. Light-sheet live imaging

Embryos were collected and glued to the outside of glass capillaries as previously described [12]. All experiments were performed at 25*^◦^*C in a Zeiss Light-Sheet Z.1 with 20*×*/1 NA Plan-Apochromat water immersion objective.

Stacks were acquired with 0.28 *µ*m XY-resolution and 2 *µ*m Z-resolution. We either imaged with 3% 561 nm laser power over half of the embryo’s surface every 1–2 minutes to reconstruct the 3D surface, or focused on the sagittal/equator section of the embryo and imaged every 15–30 seconds for subsequent PIV analysis. The balancer signal in GFP was recorded with 3% 488 nm laser power at the end of the session to check homozygosity.

### 4.4. PIV analysis

Movies were trimmed to start at gastrulation onset and finish after 40 minutes. Using Fiji [29], segmented spline fitted lines of 64px width were drawn in the dorsal area of sagittal sections between 25–75% EL. A kymograph reflecting tissue flows was obtained using the MultiKy-mograph (https://imagej.net/Multi_Kymograph) Fiji plugin. Using the straighten function, a 64 px high rectangle of this section of the embryo in time was recovered. Similarly, for equator sections, a segmented spline fitted line was drawn from the tip of the VF to the boundary of mitotic domain 3. Using the iterativePIV (https://imagej.net/plugins/piv-analyser) plugin, with 64X64px size window and masking with no threshold, displacement data in the x-axis were recovered. This displacement data was then imported into a custom pipeline in Jupyter Notebook (https://jupyter.org), where mean displacement values were extracted every 100 pixels. A quiver figure was plotted with the overlay of the original image and the displacement data converted to velocity and represented as color-coded arrows. To generate the line-plots of averaged velocity over time, the average velocity and standard deviation of each half of the blastoderm region of interest were calculated and smoothed using a Gaussian filter of σ = 1 (time-averaged).

### 4.5. Dextran injections

Oregon-R embryos were collected, mounted on their ventral side onto coverslips and allowed to dry for 15 minutes. They were covered in water and injected upon cellularization on their posterior pole with Rhodamine-B-conjugated dextran (Thermo Fisher Scientific). Dextran intensity was visualized with 1% 561 nm laser power excitation using light-sheet microscopy.

### 4.6. Laser cauterizations

Laser cauterization experiments were performed in two microscope setups: a Luxendo MuVi SPIM with a photomanipulation module for lateral cauterizations and a Zeiss LSM 780 NLO with multiphoton excitation for ventral cauterizations. For the MuVi SPIM, embryos were embedded in 2% low-melting agarose and mounted in glass capillaries to obtain *in-toto* recordings at 4 equidistant angles. A pulsed infrared laser (1030–1040 nm, 200 fs pulse duration, 1.5 W power) was used to cauterize the posterior region of the dorsal embryonic surface, attaching the blastoderm to the vitelline envelope. Using an Olympus 20*×*/1.0 NA water immersion objective, stacks with 0.29 *µ*m X-resolution and 1 *µ*m Z-resolution were acquired at four different angles every one minute. In the confocal setup, embryos were glued ventral-side down into MatTek dishes, covered in Halocarbon oil and cauterized sequentially using a near-infrared 800 nm laser (Chameleon Vision II) through a single pixel line (210 nm/px, 100 *µ*s/px), then imaged every 3 minutes for one hour with a Zeiss *×*25/0.8 NA LD LCI Plan-Apochromat glycerol immersion objective. The bottom half of the embryo was acquired with 0.21 *µ*m x-y resolution and 2 *µ*m z-step. Typically, 1 to 3 embryos were acquired simultaneously, including non-cauterized controls.

### 4.7. Landmark displacement quantifications

Relative displacement of landmarks was calculated manually in Fiji, relative to the embryo’s length, from either renderings obtained with the 3D Script plugin in Fiji [30] or sagittal sections. Angle deviation of the head (taking the anterior tip of the VF as landmark) and posterior tip of the VF were calculated from the midline in 3D ventral reconstructions using the angle measurement tool in Fiji.

### 4.8. Modeling and simulations

Our model is adapted from a previous study [15] that simulated ectopic folds in the head-trunk region driven by GB extension and the mitotic domain, with minor modifications. We simulate the entire lateral cross-section of *Drosophila* as an elastic rod with periodic boundary conditions. The total energy combines stretch and bend components:

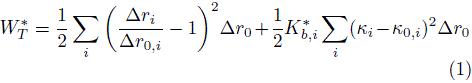

where *i* indexes each point along the rod, *r*_0,i_ is the rest length of the spring connecting consecutive points, *K_b,i_* is the dimensionless bending rigidity, and *κ_i_* and *κ*_0_*_,i_* represent the current and preferred curvatures, respectively. The value *K_b_* = 10*^−^*^4^ is adapted from [15], with modifications at invagination sites scaled by a factor of h^3^ according to experimental measurements of their relative thickness. The simulation dynamics follow the same approach as the reference model, with the addition of an adaptive time-stepping scheme that ensures no single point moves more than half the minimum inter-point distance at each step. Random noise is applied at every time step to all points except those fixed at attachment sites.

The cross-section of the embryo is modeled as an ellipse, with axis values determined from experimental measurements. At the start of the simulation, the ellipse is discretized into *N* equidistant points confined within a boundary defined by a slightly larger elliptical shape. The springs connecting neighboring points maintain a fixed distance throughout the simulation, except in the dorsal posterior region where a constriction is implemented to reflect experimental observations. In the wt case, attachment sites remain fixed; in the *scab* case, they are free to move. Invagination of the CF and PMG is modeled by setting the preferred curvature at the corresponding points to a negative value. GB extension is simulated by adding points sequentially at random locations within the GB region; after each addition, the system relaxes until spring forces fall below a threshold. When 70% of the GB points have been added, the mitotic domain points undergo doubling to represent cell division. The simulation ends once a total of 30% more points have been added to the GB and the system has relaxed.

## Acknowledgments

We thank current and former LoPaTs for discussions and support throughout this project; Anaïs Bailles, Bruno Vellutini and Fridtjof Brauns for reading and improving the manuscript with their suggestions; the Light Microscopy Facility of the Max Planck Institute of Molecular Cell Biology and Genetics (MPI-CBG) for assistance with data acquisition; Cornelia Maas for flykeeping and Sven Ssykor for his steady hand for dextran micro-injections; Yohanns Bellaïche for providing the optogenetic flies; Matt Benton for sharing HCR *scab* data; Bruno Vellutini for providing crucial fly crosses for the experiments and Abhijeet Krishna for feedback on the adaptation of the physical model.

We thank funding sources: Max Planck Society core funding to CM and PT; the European Research Council Advanced Grant (ERC-AdG 885504 GHOSTINTHESHELL) to PT; financial support from the Volkswagen Foundation under the Life initiative (AZ 98218) to WYY; the Max Planck POSTECH/Korea Research Initiative Internship Program to YJK and open access funding provided by Max Planck Society.

## Author Contributions

MBC, CM and PT conceived the study. MBC kept fly stocks, designed, performed and analyzed all the live-imaging experiments and performed the pHist-3 immunostaining. GS performed fly crosses, conducted and quantified the HCR, and the optogenetic experiments. WYY and YJK adapted and extended the model, and WYY performed the simulations and corresponding plots. MBC, PT and CM drafted the manuscript. All authors contributed to and revised the manuscript.

## Data Availability

Supplementary videos can be found in Zenodo. The code for the physical model simulations is available on gitlab. Raw data and PIV post-processing code is available on Zenodo and github.

---

**Figure S1.**
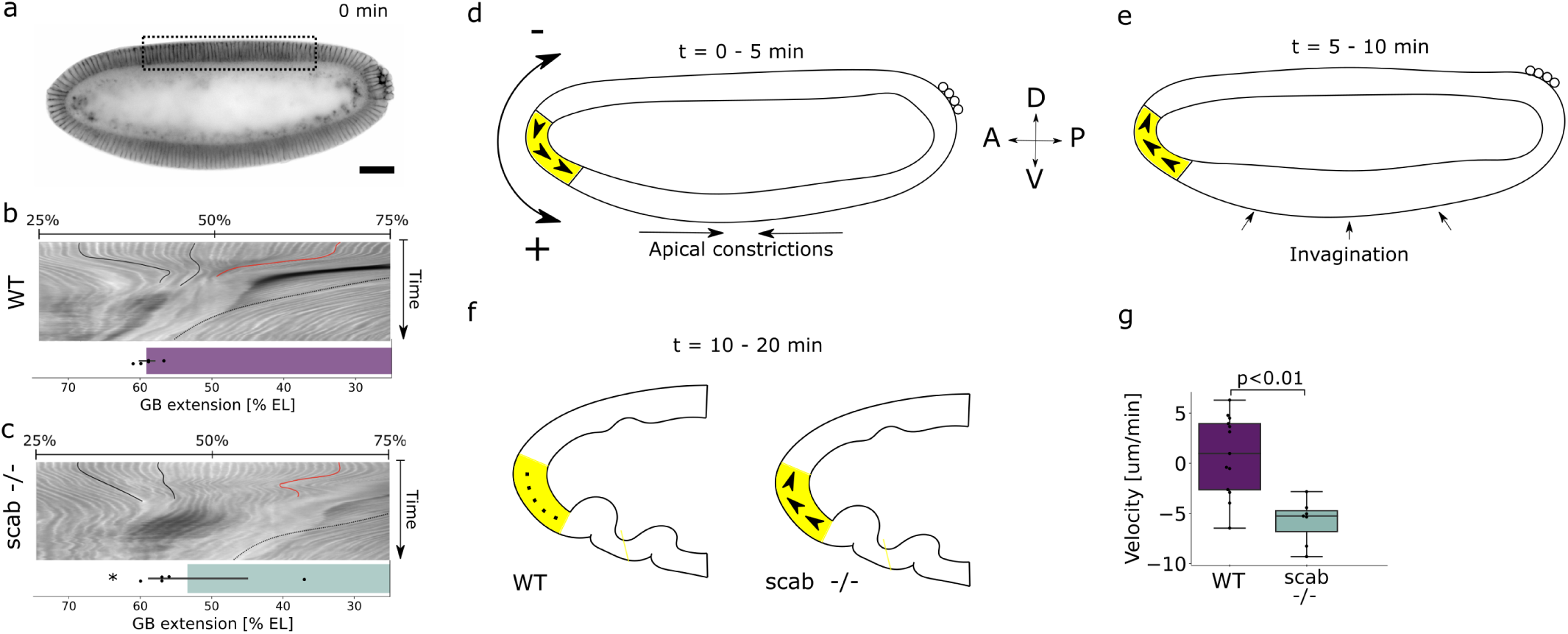
(a) Sagittal section of Dorophila embryo at timepoint 0 and region where kymographs were extracted from. (b, c) Kymograph of representative wt (b) and scab mutant embryos with cephalic furrow invagination area (solid black lines), GB extension (dotted black line) and tissue displacement (solid red line) highlighted. Barplot of average GB extension, 95% CI and single data points (N=5). (d) Counter-clockwise tissue flows in the anterior pole during early gastrulation (0-5 min). (e) Clockwise tissue flows during VF invagination (5-10 min). (f) Tissue flows stop in wt embryos and continue in scab mutants (10-20 min). (g) Boxplot of quantified tissue flows in figure 1d at t = 15 min and individual data points. Boxes indicate the quartiles and the bars the full extension of the distribution. Scale bar equals 50 um.

**Figure S2.**
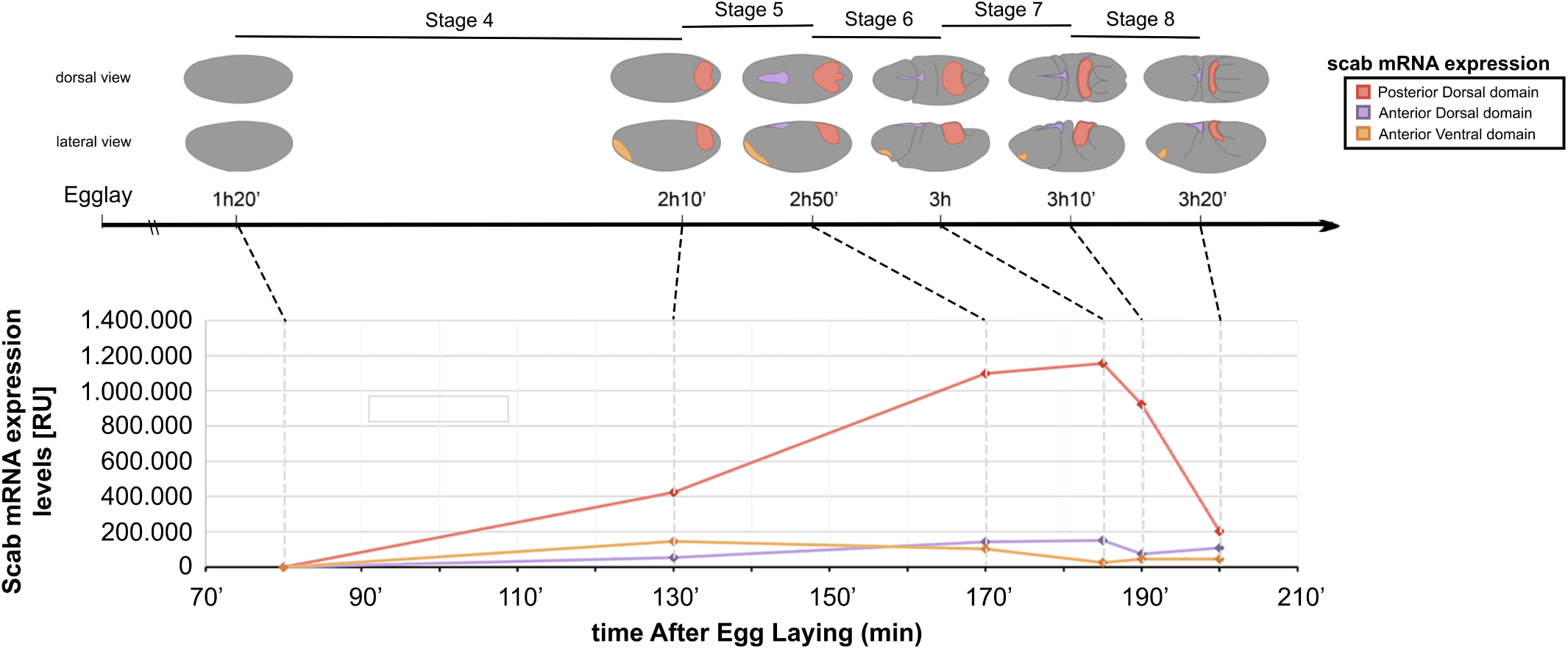
Diagram of embryo morphology at different embryonic stages and regions of Scab expression. Quantified HCR signal of each domain is plotted for each stage to observe the dynamics of Scab expression.

**Figure S3.**
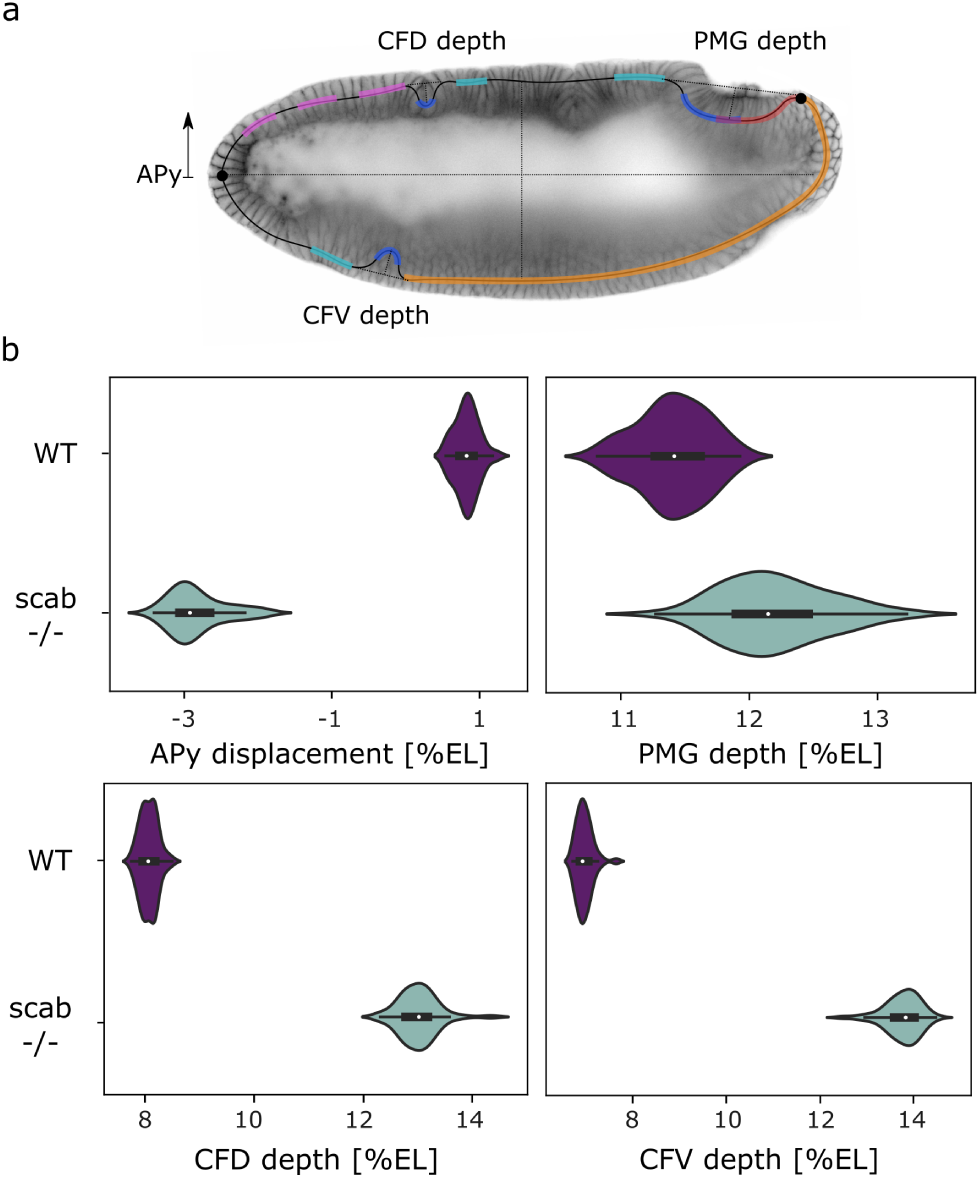
(a) Additional quantified features (APy displacement, PMG, CFD and CFV depths). (b) Violin plots of the distributions of the quantified features for the attachments ON (wt scenario) and OFF (*scab* scenario).

**Figure S4.**
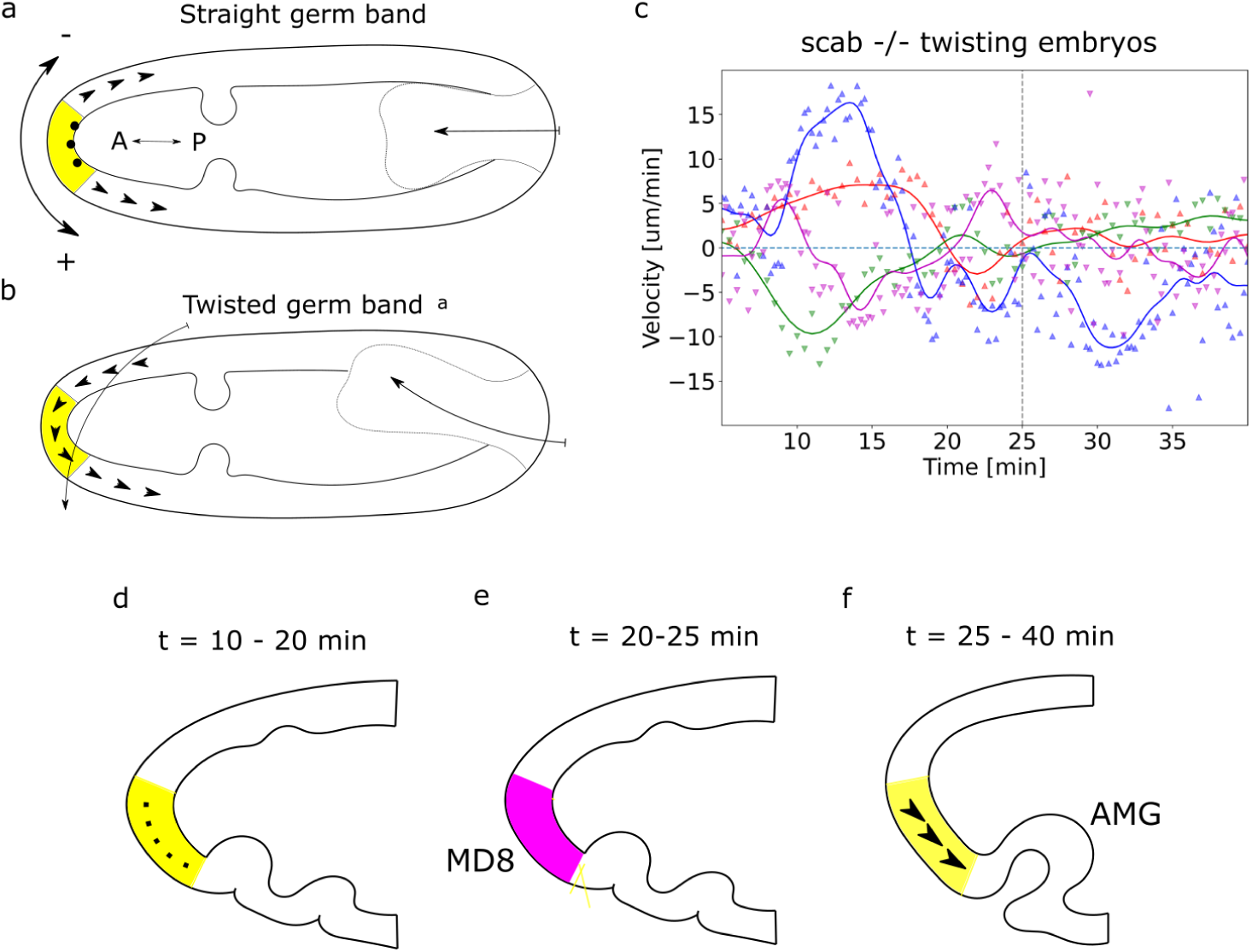
(a, b) Diagram of tissue flows on the anterior pole in the equator sections of embryos with a straight GB extension versus twisting GB. (c) Single curves of average velocities in anterior pole equator sections for twisting scab mutants (N = 4) showing non-stereotypical behavior.

**Figure S5.**
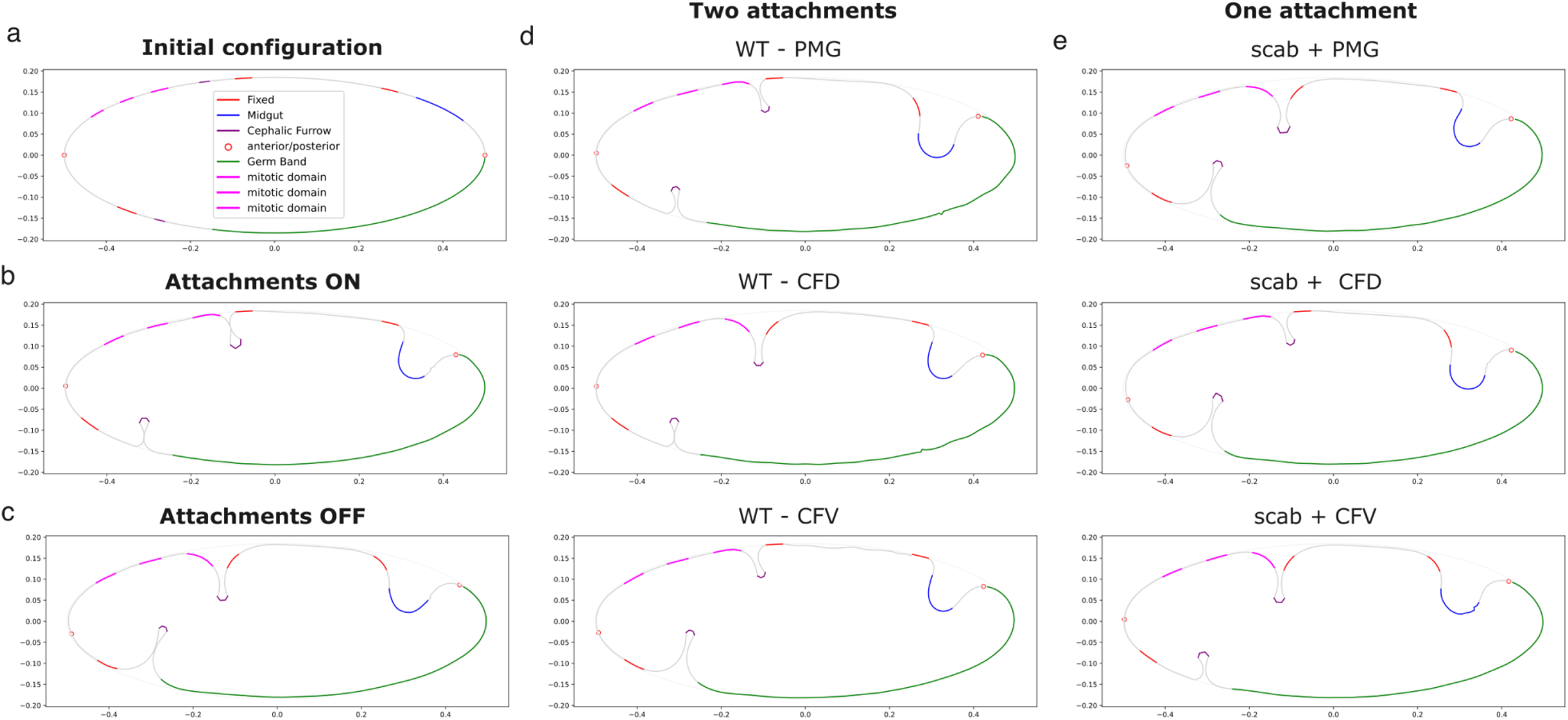
(a) Initial configuration in simulated curved embryo surface with highlighted regions. (b) Final simulation results for a representative all OFF (*scab*) and all ON (wt) situation. (c) Final simulation results for a representative of two attachment simulations. (c) Final simulation results for a representative single attachment simulation.

**Figure S6.**
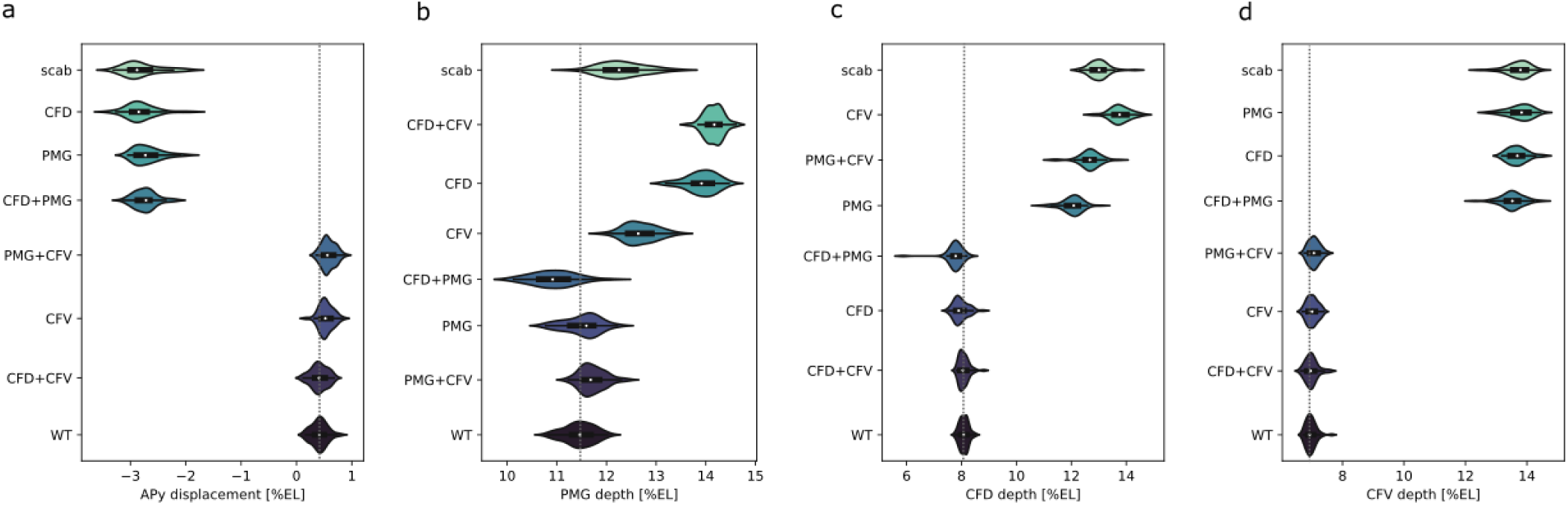
(a, b, c, d) Violin plots of the additional quantified features’ locations ranked by relevance in the single and double attachment situations compared to all attachments ON (wt) and all OFF (*scab*). The gray dotted line highlights the mean of the wt situation.

## References

[1] Raphaël Clément, Benoît Dehapiot, Claudio Collinet, Thomas Lecuit, and Pierre-François Lenne. Viscoelastic dissipation stabilizes cell shape changes during tissue morphogenesis. Current biology, 27(20):3132– 3142, 2017.

[2] Diana Khoromskaia and Guillaume Salbreux. Active morphogenesis of patterned epithelial shells. eLife, 12:e75878, January 2023.

[3] Jana F. Fuhrmann, Vincenzo Maria Schimmenti, Greta Cwikla, Sangwon Lee, Michaela Yuan, Michaela Wilsch-Bräuninger, Frank Jülicher, Marko Popović, and Natalie A. Dye. Apical extracellular matrix regulates fold morphogenesis in the Drosophila wing disc, October 2025. ISSN: 2692-8205 Pages: 2025.09.06.674631 Section: New Results.

[4] T. Pascucci, J. Perrino, A. P. Mahowald, and G. L. Waring. Eggshell assembly in Drosophila: processing and localization of vitelline membrane and chorion proteins. Developmental Biology, 177(2):590–598, August 1996.

[5] Stefan Münster, Akanksha Jain, Alexander Mietke, Anastasios Pavlopoulos, Stephan Grill, and Pavel Tomancak. Attachment of the blastoderm to the vitelline envelope affects gastrulation of insects. Nature, 568, April 2019.

[6] Anaïs Bailles, Claudio Collinet, Jean-Marc Philippe, Pierre-François Lenne, Edwin Munro, and Thomas Lecuit. Genetic induction and mechanochemical propagation of a morphogenetic wave. Nature, 572:1–7, August 2019.

[7] Claudio Collinet, Anaïs Bailles, Benoit Dehapiot, and Thomas Lecuit. Mechanical regulation of substrate adhesion and de-adhesion drives a cell-contractile wave during *Drosophila* tissue morphogenesis. Developmental Cell, 59(1):156–172.e7, January 2024.

[8] Serafini, Maryam Setoudeh, Marina B. Cuenca, Charlène Brillard, Matthias Arzt, Pavel Mejstřik, Pierre A. Haas, and Pavel Tomančák. Embryo-eggshell interaction counteracts chiral bias in early Drosophila morphogenesis. bioRxiv, March 2026.

[9] Annick Sawala, Margherita Scarcia, Catherine Sutcliffe, Scott G. Wilcockson, and Hilary L. Ashe. Peak BMP Responses in the Drosophila Embryo Are Dependent on the Activation of Integrin Signaling. Cell Reports, 12(10):1584–1593, 2015.

[10] Celia M. Smits, Sayantan Dutta, Vishank Jain-Sharma, Sebastian J. Streichan, and Stanislav Y. Shvartsman. Maintaining symmetry during body axis elongation. Current Biology, 33(16):3536–3543.e6, August 2023.

[11] Peter A. Santi. Light Sheet Fluorescence Microscopy. Journal of Histochemistry and Cytochemistry, 59(2):129–138, February 2011.

[12] Christopher Schmied and Pavel Tomancak. Sample Preparation and Mounting of Drosophila Embryos for Multiview Light Sheet Microscopy. In Christian Dahmann, editor, Drosophila: Methods and Protocols, pages 189–202. Springer New York, New York, NY, 2016.

[13] Matteo Rauzi, Uros Krzic, Timothy E. Saunders, Matej Krajnc, Primoz Ziherl, Lars Hufnagel, and Maria Leptin. Embryo-scale tissue mechanics during Drosophila gastrulation movements. Nature Communications, 6, 2015.

[14] A. Vincent, J. T. Blankenship, and E. Wieschaus. Integration of the head and trunk segmentation systems controls cephalic furrow formation in Drosophila. Development, 124(19):3747–3754, October 1997.

[15] Bruno C. Vellutini, Marina B. Cuenca, Abhijeet Krishna, Alicja Szalapak, Carl D. Modes, and Pavel Tomancak. Patterned invagination prevents mechanical instability during gastrulation. Nature, 646(8085):627–636, October 2025.

[16] Anna Popkova, Urška Andrenšek, Sophie Pagnotta, Primož Ziherl, Matej Krajnc, and Matteo Rauzi. A mechanical wave travels along a genetic guide to drive the formation of an epithelial furrow during *Drosophila* gastrulation. Developmental Cell, 59(3):400–414.e5, February 2024.

[17] Bipasha Dey, Verena Kaul, Girish Kale, Maily Scorcelletti, Michiko Takeda, Yu-Chiun Wang, and Steffen Lemke. Divergent evolutionary strategies pre-empt tissue collision in gastrulation. Nature, 646(8085):637– 646, October 2025.

[18] Dominik Stappert, Nadine Frey, Cornelia von Levetzow, and Siegfried Roth. Genome-wide identification of Tribolium dorsoventral patterning genes. Development, 143(13):2443–2454, July 2016.

[19] Victoria E. Foe. Mitotic domains reveal early commitment of cells in Drosophila embryos. Development, 107(1):1–22, September 1989. eprint: https://journals.biologists.com/dev/article-pdf/107/1/1/2794782/develop_107_1_1.pdf.

[20] Florencia di Pietro, Sophie Herszterg, Anqi Huang, Floris Bosveld, Cyrille Alexandre, Lucas Sancéré, Stéphane Pelletier, Amina Joudat, Varun Kapoor, Jean-Paul Vincent, and Yohanns Bellaïche. Rapid and robust optogenetic control of gene expression in Drosophila. Developmental Cell, 56(24):3393–3404.e7, December 2021.

[21] Mahamar Dicko, Pierre Saramito, Guy B. Blanchard, Claire M. Lye, Bénédicte Sanson, and Jocelyn Étienne. Geometry can provide long-range mechanical guidance for embryogenesis. PLOS Computational Biology, 13(3):e1005443, March 2017.

[22] Paul Z. Liu and Thomas C. Kaufman. Short and long germ segmentation: unanswered questions in the evolution of a developmental mode. Evolution & Development, 7(6):629–646, 2005. eprint: https://onlinelibrary.wiley.com/doi/pdf/10.1111/j.1525-142X.2005.05066.x.

[23] Jeremy A. Lynch, Ezzat El-Sherif, and Susan J. Brown. Comparisons of the embryonic development of Drosophila, Nasonia, and Tribolium. WIREs Developmental Biology, 1(1):16–39, 2012. eprint: https://wires.onlinelibrary.wiley.com/doi/pdf/10.1002/wdev.3.

[24] R. Murakami, S. Takashima, and T. Hamaguchi. Developmental genetics of the Drosophila gut: specification of primordia, subdivision and overt-differentiation. Cellular and Molecular Biology, 45(5):661–676, July 1999.

[25] Steffen Lemke, Girish Kale, and Silvia Urbansky. Comparing gastrulation in flies: Links between cell biology and the evolution of embryonic morphogenesis. Mechanisms of Development, 164:103648, December 2020.

[26] Reinhard Schröder, Christoph Eckert, Christian Wolff, and Diethard Tautz. Conserved and divergent aspects of terminal patterning in the beetle Tribolium castaneum. Proceedings of the National Academy of Sciences of the United States of America, 97(12):6591–6596, June 2000.

[27] Nicola Berns, Thomas Kusch, Reinhard Schröder, and Rolf Reuter. Expression, function and regulation of Brachyenteron in the short germband insect Tribolium castaneum. Development Genes and Evolution, 218(3):169–179, April 2008.

[28] Harry M. T. Choi, Maayan Schwarzkopf, Mark E. Fornace, Aneesha Acharya, Georgios Artavanis, J. Stegmaier, Alexandre Cunha, and Niles A. Pierce. Third-generation in situ hybridization chain reaction: multiplexed, quantitative, sensitive, versatile, robust. Development, 145, 2018.

[29] Curtis T. Rueden, Johannes E. Schindelin, Mark C. Hiner, Barry E. DeZonia, Alison E. Walter, Ellen T. Arena, and Kevin W. Eliceiri. ImageJ2: ImageJ for the next generation of scientific image data. BMC Bioinformatics, 18, 2017.

[30] Benjamin Schmid, Tina Fraass, Philipp Tripal, Jan Huisken, Christina Kersten, and Ralf Palmisano. 3Dscript: Animating 3D/4D microscopy data using a natural language based syntax. Nature Methods, 16, March 2019.

